# VESTIBULAR FUNCTION LOSS ASSOCIATES WITH SENSORY EPITHELIUM PATHOLOGY IN VESTIBULAR SCHWANNOMA PATIENTS

**DOI:** 10.64898/2026.03.23.713132

**Authors:** Mireia Borrajo, Àngela Callejo, Elisabeth Castellanos, Emilio Amilibia, Jordi Llorens

**Affiliations:** Departament de Ciències Fisiològiques, Institut de Neurociències, Universitat de Barcelona (UB), Feixa Llarga s/n, 08907 l’Hospitalet de Llobregat, Catalunya, Spain; Institut d’Investigació Biomèdica de Bellvitge (IDIBELL), 08907 l’Hospitalet de Llobregat, Catalunya, Spain; Servei d’Otorinolaringologia, Hospital Universitari Germans Trias i Pujol, Badalona, Catalunya, Spain; Servei de Genètica, LCMN, Hospital Universitari Germans Trias i Pujol, Badalona, Catalunya, Spain; Clinical Genomics Research Group, Institut de Recerca Germans Trias i Pujol (IGTP), Badalona, Catalunya, Spain

**Keywords:** Vestibular schwannoma, vestibular hair cells, NF2 gene, vestibulo-ocular reflex, calyx afferents

## Abstract

Vestibular schwannomas (VS) cause vestibular function loss by mechanisms still poorly understood. We evaluated the vestibulo-ocular reflex by the video-assisted Head Impulse Test (vHIT) in patients with planned tumour resection by a trans-labyrinthine approach. The vestibular sensory epithelia were collected and processed by immunofluorescent labelling for confocal microscopy analysis of sensory hair cell subtypes (type I, HCI, and type II, HCII), calyx endings of the pure-calyx afferents, and the calyceal junction normally found between HCI and the calyx (n=23). Comparing Normo-vHIT and Hypo-vHIT patients, we concluded that worse vestibular function associates with decreased HCI and HCII counts in the sensory epithelia and with increased proportion of damaged calyces. A decrease in the number of HCI and calyx endings of the pure-calyx afferents was recorded to associate with age increase. Partial least squares regression (PLSR) models indicated that VS and age had independent, additive effects on vestibular function. Correlation analyses indicated that lower vHIT gains associate with lower numbers of HCI and increased percentages of damaged calyces. These data support the hypothesis that the deleterious effect of VS on vestibular function is mediated, at least in part, by its damaging impact on the vestibular sensory epithelium. They also provide further evidence for the dependency of the vestibulo-ocular reflex on HCI function and for the calyceal junction pathology as a common response of the sensory epithelium to HC stress.

## INTRODUCTION

The vestibular system encodes information on the movements of the head and body within the gravitational field of the earth through five sensory epithelia in each ear: three cristae in the semicircular canals and two otolith receptors, the utricle and the saccule. The sensory epithelia of these five vestibular receptors have similar composition, comprised of a pseudostratified epithelium made of supporting cells and two types of transducing hair cells (HCs), type I (HCI) and type II (HCII). HCI and HCII have several morphological, molecular and functional differences, with HCI sustaining very fast transmission of vestibular signals (Eatock and Songer, 2011; Govindaraju et al., 2023; McInturff et al., 2018). HCI are encased by a unique kind of afferent terminal, named calyx, which forms a specialized adhesion complex, the calyceal junction (Lysakowski et al., 2011; Sedó-Cabezón et al., 2015), that maintains a narrow spacing between the pre- and post-synaptic sides that is necessary for the fast non quantal transmission that characterizes the unit (Govindaraju et al., 2023). The central part of the receptors, known as striola in the utricle and saccule, contains a subset of afferents, named pure-calyx or calyx-only afferents, that express the calcium binding protein calretinin (Dechesne et al., 1991; Lysakowski et al., 2011).

Like other sensory systems, the vestibular system shows an age-dependent loss of function and sensory receptor decline (Baloh et al., 2001; Velázquez-Villaseñor et al., 2000; Zalewski, 2015). In addition, many different insults, including vestibular schwannoma (VS) and exposure to ototoxic chemicals, can trigger vestibular function loss. VS are Schwann cell-derived tumours that grow on the vestibulocochlear nerve (Carlson and Link, 2021). VS is the most frequent tumour of the cerebellopontine angle and constitute about 8 % of intracranial tumours. Although benign, VS causes significant morbidity, including sensorineural hearing loss in > 90% and vestibular disturbances in > 60 % of patients (Carlson and Link, 2021; Matthies and Samii, 1997). Most cases (>95%) of VS are sporadic and unilateral but others are due to genetic syndromes. Thus, almost all patients affected by NF2-related schwannomatosis (NF2-SWN, previously known as neurofibromatosis type 2) develop bilateral VS (Plotkin et al., 2022). Most sporadic VS also have loss-of-function mutations of the NF2 (NF2, moesin-ezrin-radixin like [MERLIN] tumour suppressor) gene, and it has been demonstrated that the NF2 disruption is a key event in VS development (Jacoby et al., 1996; Vitte et al., 2017).

The pathological mechanisms responsible for both age-associated and VS-associated vestibular function loss remain insufficiently understood. A deeper understanding of these mechanisms may guide the development of prophylactic approaches to preserve function or may open the use of functional data to infer tumour characteristics. In VS, it was initially assumed that mechanical compression of the nerve would cause the functional deficits (Nadol et al., 1996; Roosli et al., 2012), but several observations, including the lack of correlation between the radiographic tumour size and the audiometric thresholds shifts, revealed that other mechanisms may be involved (Caye-Thomasen et al., 2007; Nadol et al., 1996). To address this question, audiovestibular nerves and sensory epithelia from the inner ear of VS patients have been studied through post-mortem processing of temporal bones (Brown et al., 2025; Hızlı et al., 2016) or through isolation during excision of the tumour by the trans-labyrinthine approach (Sans et al., 1996; Wang et al., 2024), a major therapeutic option when tumour growth becomes life-threatening. Studies using either approach have revealed multiple mechanisms involved in VS-induced audiovestibular dysfunction. Thus, there are reports of cases in which the growth of the tumour extending inside the inner ear resulted in direct damage to the organ of Corti or the vestibular sensory epithelium (Brown et al., 2025; Ylikoski et al., 1978). However, the symptoms typically appear before inner ear invasion, consequent to a damaging effect of the tumour on the audiovestibular nerve (Benitez et al., 1967; Sans et al., 1996; Ylikoski et al., 1978) or to loss of HCs (Hızlı et al., 2016; Sans et al., 1996; Wang et al., 2024). In the organ of Corti, increasing evidence supports the conclusions that the extent of hearing loss may relate to tumour biology through diffusible factors secreted by the tumour that have a deleterious impact on the HC (Dilwali et al., 2015; Sagers et al., 2019; Soares et al., 2016; Vasilijic et al., 2025). By contrast, much less attention has been placed on the vestibular system, even though imbalance is more disabling than hearing loss in these patients (Lloyd et al., 2010).

In animal models of chronic ototoxicity using the experimental compound 3,3’-iminodipropionitrile, dismantlement of the calyceal junction and parallel synaptic uncoupling represent early pathological events associated with the initial functional loss at the organism level (Greguske et al., 2019; Sedó-Cabezón et al., 2015). A prominent feature is the loss of the adhesion protein that characterizes the junction, contactin-associated protein (CASPR1) (Sousa et al., 2009). Interestingly, this initial pathology is reversible and the histological repair associates with functional recovery (Greguske et al., 2019; Sedó-Cabezón et al., 2015). More recently, dismantlement of the calyceal junction was observed in vestibular epithelia of rats exposed to the clinically-relevant ototoxic aminoglycoside antibiotic, streptomycin, and of VS patients, leading to the hypothesis that HC-calyx detachment is a common response of the vestibular epithelium under stress of diverse origin (Maroto et al., 2023). Therefore, the relevance of calyceal junction dismantlement in VS merited a detailed study. Differential sensitivity to ototoxic agents and to age-related decline is also a known feature distinguishing the two types of HCs, being HCI the first ones to be lost (Anniko, 1983; Maroto et al., 2023, 2021; Zalewski, 2015). In addition, HCI synapsed by pure-calyx afferents are the most sensitive to ototoxic damage (Maroto et al., 2021; Schenberg et al., 2026).

The aim of this study has been to correlate vestibular function and sensory epithelium damage in VS patients to reveal pathological events mediating the function loss in VS. Among the options for vestibular function assessment, we selected the video-head impulse test (vHIT) for semicircular canal function (Weber et al., 2009) and the vestibular evoked myogenic potentials (VEMPS) for otolith function (Rosengren et al., 2019). While these tests do not cover the full range of vestibular functions, they are non-invasive, quick and reliable test that cause little discomfort to the patients. Also, VS patients requiring trans-labyrinthine tumour excision can provide sensory epithelia for molecular, cell or tissue studies (Taylor et al., 2018, 2015; Wang et al., 2024). Although at least one study has used these epithelia to assess the integrity of the sensory epithelium by state-of-the-art immunofluorescent analysis (Wang et al., 2024), to our knowledge no previous work has correlated such analysis to the pre-surgery vestibular function of the patients. Therefore, the histological integrity of the vestibular epithelium regarding calyceal junctions, afferent terminals, HCI and HCII was evaluated here in relation to vHIT responses and age in samples of VS patients.

## MATERIALS AND METHODS

### Subjects

This study included 23 patients presenting unilateral VS confirmed by magnetic resonance imaging (MRI) using T1- and T2-weighted sequences with high-resolution (0.5 mm) slices through the internal auditory canal, who underwent surgical resection of the tumour at the Otorhinolaryngology Department, Hospital Universitari Germans Trias i Pujol (HUGTP) (Badalona, Catalunya, Spain) in the period July 2020 to June 2023, and follow up at Spanish Center of Phakomatoses (CSUR). Seventy percent were male and the mean ± SD age was 50.4 ± 15.7 years. The concomitance confounding conditions such as Ménière’
ss disease, immunomediated inner ear disease, ototoxic drug intake, inner ear malformations, or tumour invasion of the inner ear, were considered excluding criteria. Patients in which the histological evaluation failed to provide useful results were also excluded from the study. This study adhered to the tenets of the Declaration of Helsinki and was approved by the hospital’s Clinical Ethics Committee (PI-18-166). Informed consent was obtained from all individual participants included in the study.

### Vestibular functional study

Semicircular canal function was evaluated using an ICS impulse three-dimensional vHIT system (GN Otometrics, Taastrup, Denmark). All the examinations were performed by the same explorer. Both horizontal (lateral) and vertical (anterior and posterior) canals were examined bilaterally. Eye and head velocities were sampled at 250 Hz and vestibulo-ocular reflex (VOR) gain (ratio of eye-to-head velocity) was calculated from the average of at least 10 responses to head impulses performed over a range of velocities (50-300º/s). Normal gain was defined as ≥0.80 in the horizontal canals and ≥0.70 in the vertical canals (McGarvie et al., 2015), and patients with a lower value in any of the canals was classed in the Hypo-vHIT group. Also gain asymmetry according to Newman-Toker formula (Zamaro et al., 2020) was calculated as a relative value for gain. Presence of catch-up saccades (CUS), their appearance in relation to the head movement and their organization, were taken into consideration (Rey-Martinez et al., 2015). No severe adverse events were associated with the realization of the test. In this series of patients, VEMPS were also assessed as described elsewhere (Callejo et al., in preparation). However, only 6 of the 23 patients for which histological data became available had measurable VEMP results, so these data were excluded from the study.

### Immunofluorescent analysis of the vestibular epithelium

SV resection was performed by the translabyrinthine surgical approach. The vestibular epithelia were harvested and immediately placed in 4% formaldehyde with < 2.5% methanol (BiopSafe). They were kept at 4ºC and sent to the laboratory the next day, except for two samples that were delayed till day 4. As reported previously (Maroto et al., 2023), this delay has no impact on the histological results. Upon arrival, the vestibular epithelia were cleaned from adjacent tissues, rinsed in phosphate buffered saline (PBS, pH 7.2, 2 times 20’), transferred into a cryoprotective solution (34.5% glycerol, 30% ethylene glycol, 20% PBS, 15.5% distilled water), allowed 2 h of embedding at 4ºC and stored at – 20 °C until further processing.

Immunohistochemical fluorescence staining and confocal microscopy analysis were performed as described in detail elsewhere (Maroto et al., 2023). Briefly, tissues were rinsed with PBS to remove the cryoprotective solution, then immunolabelled with rabbit anti-Myosin VII a (MYO7A, 25-6790, Proteus Biosciences, RRID: AB_10015251; used at 1:400 dilution), mouse anti CASPR1 (clone K65/35, RRID: AB_2083496; 1:400), and guinea-pig anti-calretinin (214.104, Synaptic Systems, RRID: AB_10635160; 1:500) antibodies. Alexa-405, -488, - 555, and -647-conjugated secondary antibodies (Invitrogen/Thermo Fisher) were used to reveal the primary antibodies. Whole-mount preparations were imaged in a Zeiss LSM880 spectral confocal microscope. Two images were obtained using a 40X objective (numeric aperture 1.3), one from the striolar/central area of the receptor and one from the peripheral area. The images were quantified manually using the cell counter tool of the Image J software (National Institute of Mental Health, Bethesda, Maryland, USA).

### Genetic NF2 test

When possible, Schwann cells from vestibular schwannomas were cultured as described previously (Catasús et al., 2025) and DNA from the VS or cell culture was tested for NF2 variants using the customized I2HCP panel (Castellanos et al., 2017). All identified variants were annotated using NM_000268.3 and hg17 as reference sequences.

### Statistics

Raw histological data were compared between groups of sex and vestibular function with the Mann-Whitney U test. The relationships between age and the histological variables were evaluated using Pearson’s correlation tests. These analyses were performed using the SPSS Statistics 29 program package. To study the interaction between the function and age factors, we elaborated partial least squares regression (PLSR) models, using the R-Studio programme package. These models provide a good approach to multicollinearity and non-normality of the variables under study (Lafaye De Micheaux et al., 2019). They were validated with the jack-knife estimation of parameter uncertainty (Martens and Martens, 2000). Then, regression coefficients were obtained from the jack.test R function. The number of components was selected using the summary function of the PLSR package to minimize the error of the model and maximize the percentage of variability explained. Finally, we used the Pearson’s correlation test to assess the relationship between the histological measures and the data from the vHIT tests from the tumour side and the contralateral side. The Holm correction was used to adjust the p-values for multiple tests (Xie, 2012). Significance level was set at p<0.05.

## RESULTS

The 23 patients were sorted in two groups according to their vHIT responses, those with preserved responses (Normo-vHIT, n=11) and those with impaired responses (Hypo-vHIT, n=12). In 18 of the cases, the genetic analysis of the tumour was available. As shown in Supplementary Table 1, all cases exhibited genetic alterations that impaired the function of the NF2 gene, including splicing, stop codon and frameshift mutations, as described previously in sporadic and NF2-related scwhannomatosis (Catasús et al., 2025; Evans et al., 2020; Halliday et al., 2019, 2017; Jacoby et al., 1996). Nine cases showed complete NF2 loss: four cases had two different point mutations in the NF2 gene; four exhibited a variant frequency higher than 50%, indicating loss of heterozygosity (LOH) of the other NF2 allele; and one case showed the deletion of both chromosome 22. In the other nine cases, a single mutation was found, denoting the inactivation of one copy of the gene. Due to contamination by non-tumoral cells, it is also possible that they had loss of the wild-type allele, which was undetectable in this study.

Confocal microscopy analysis of the samples (Fig. 1) allowed the identification of HCs by their MYO7A expression. The calyces encasing HCI were identified by CASPR1 label. This label characterizes calyces in normal rodent tissues (Lysakowski et al., 2011; Sedó-Cabezón et al., 2015; Sousa et al., 2009) and has been reported with identical features in samples of several non-VS human samples (Maroto et al., 2023). Calretinin expression identified HCII and, in the central/striola regions, also the calyx terminals of the pure-calyx afferents. These pure-calyx/calretinin+ afferents are not present in the periphery of the receptors. In the normal rodent and non-SV human samples, the calretinin+ afferents are also CASPR1+ (Lysakowski et al., 2011; Maroto et al., 2023), so lack of CASPR1 in calretinin+ calyces in the of SV patients denoted absence of the normal calyceal junction and these calyces were therefore counted as damaged calyces.

**Figure 1.**
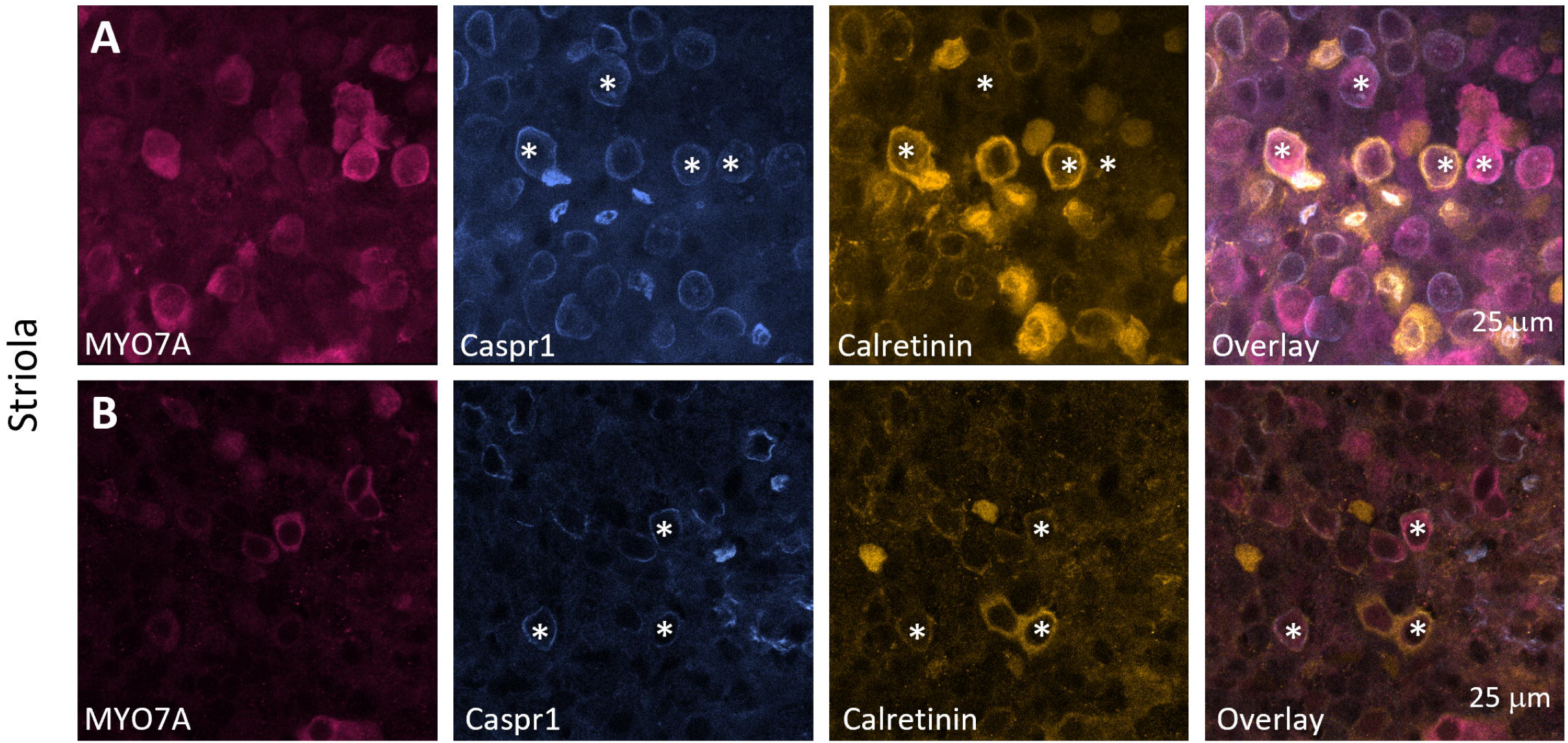
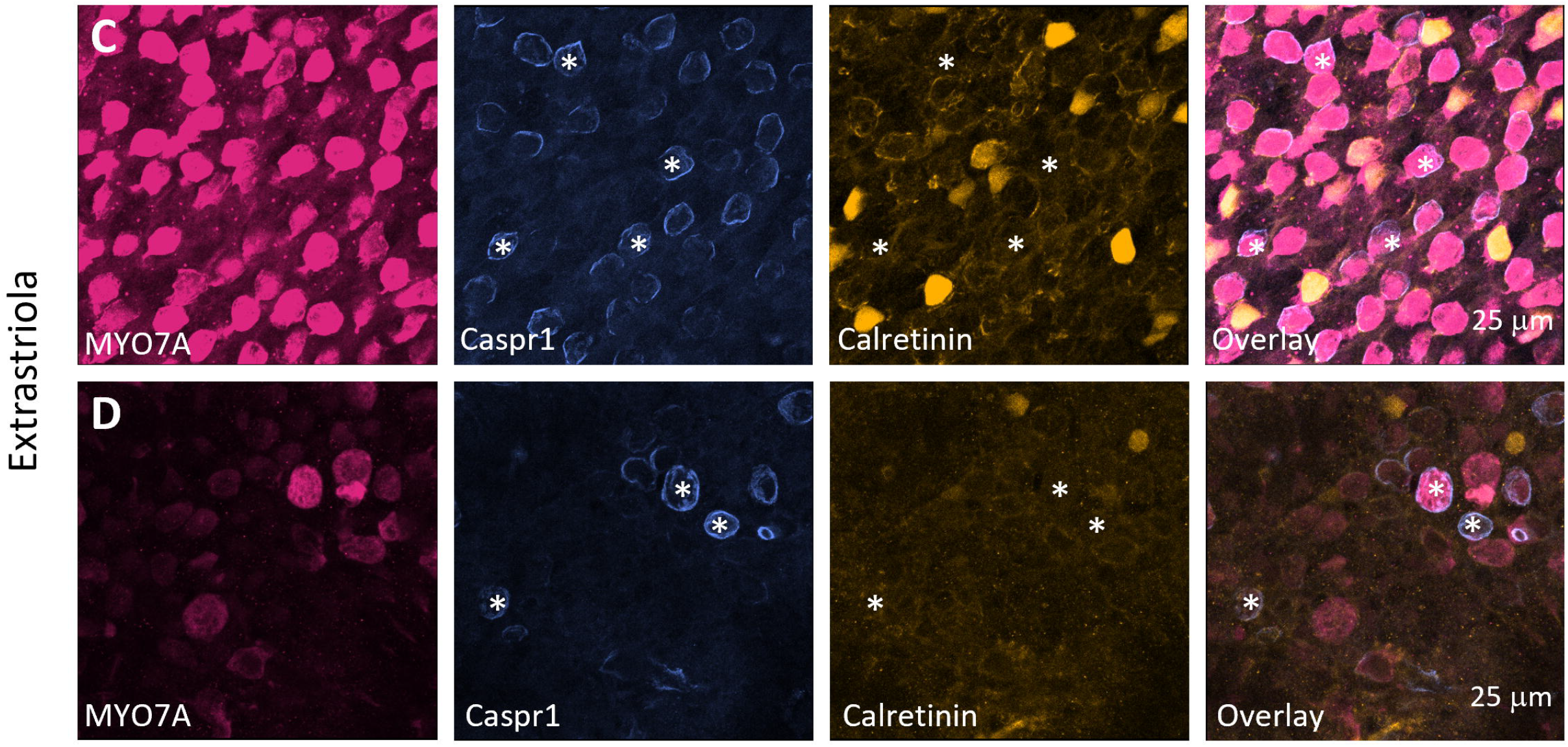
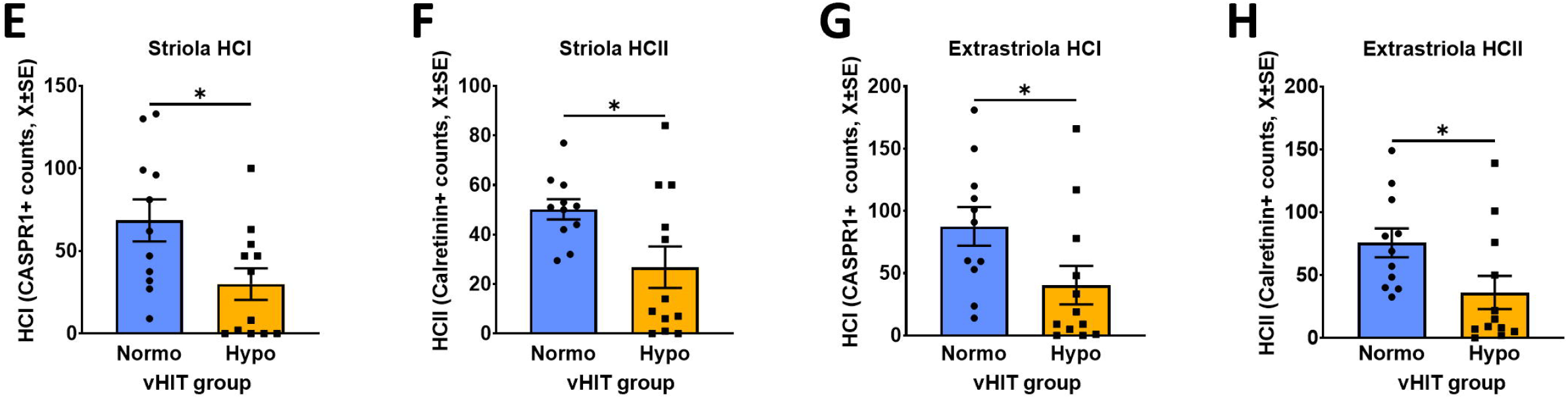
Density of HCI and HCII in the vestibular sensory epithelium according to epithelium area (striola vs. periphery) and functional group (Normo-vHIT vs. Hypo-vHIT) in vestibular schwannoma (VS) patients. **A-D:** Confocal microscopy images showing immunolabel for MYO7A (magenta, left panels), CASPR1 (blue, second column), and calretinin (yellow, third column). The right-most column shows the overlay of the three labels. Rows **A** and **B** are from the striola region and rows **C** and **D** are from the periphery of patients from the Normo-vHIT group (**A** and **C**) and the Hypo-vHIT group (**B** and **D**). Annotations mark CASPR1+ profiles identifying HCI (asterisks), calretinin+ profiles identifying HCII (arrows) and calretinin profiles identifying pure-calyx afferents (arrowheads). **E-H:** Quantitative analysis. The graphs show individual values and mean ± SEM of HCI (**E** and **G**) and HCII (**F** and **H**) counts in the striola (**E** and **F**) and periphery (**G** and **H**) regions. *: p< 0.05, Mann-Whitney U test.

Comparison of the histological variables between sexes did not find any significant difference in any of the variables (data not shown) so data from male and female patients were pooled together for subsequent analyses. Type specific HC assessment revealed statistically significant group differences. Thus, Hypo-vHIT patients had reduced counts of HCI (CASPR1+) and HCII (calretinin+) in both the central (Fig. 1 A, B, E, F) and peripheral (Fig. 1. C, D, G, H) regions of the sensory epithelium. Mann-Whitney U and p values were, respectively, as follows: 32 and 0.041 (HCI in striola); 28 and 0.018 (HCI in periphery); 33 and 0.043 (HCII in striola); and 28 and 0.019 (HCII in periphery). In addition, patients of the Hypo-vHIT group had a highly significant (U=13, p=0.0005) increase in the percentage of calyces from pure-calyx afferents that were damaged, that is, an increased proportion of calretinin+ calyces devoid of the normal CASPR1 label (Fig. 2).

**Figure 2.**
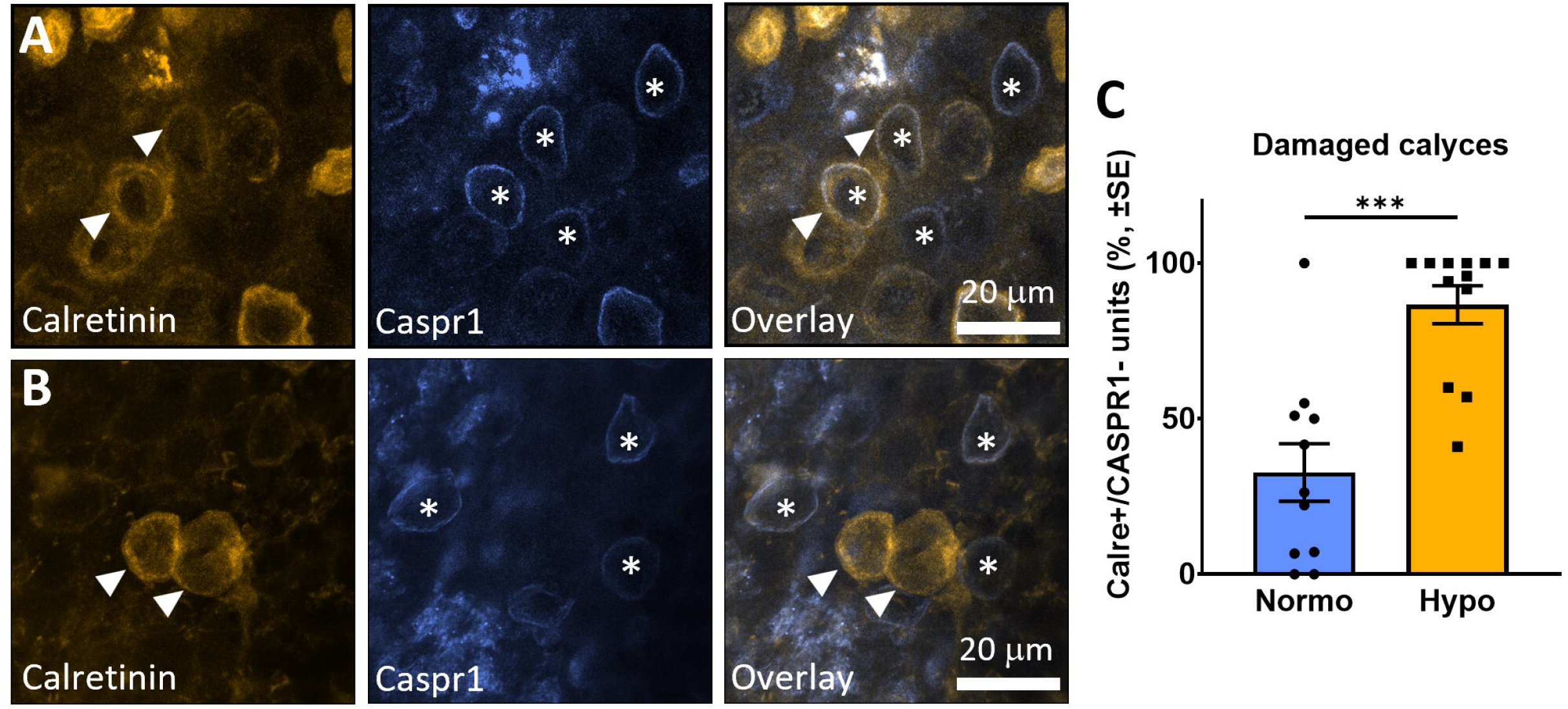
Detachment of HCI from pure-calyx afferents in the vestibular epithelium and its association with vestibular function. **A-B:** Confocal microscopy images showing immunolabel for calretinin (yellow, left panels), CASPR1 (blue, middle panels), and both calretinin and CASPR1 (left panels) of a patient from the Normo-vHIT group (**A**) and one from the Hypo-vHIT group (**B**). In **A**, the calretinin+/pure-calix afferents (arrowheads) have a calyceal junction identified by the CASPR1+ label (asterisks). CASPR1+ calyces without calretinin label are also observed. In **B**, note two calretinin+/pure-calix afferents with no CASPR1 label, an abnormal feature of the epithelium. (**C**) Group comparison (Normo-vHIT vs. Hypo-vHIT) of the percentage of calretinin+ calyces (pure-calyx afferents) with abnormal lack of CASPR1 label. ***: p< 0.001, Mann-Whitney U test.

The differences among groups were not a consequence of a bias due to age of the patients, because both Normo-vHIT and Hypo-vHIT groups included a wide range of ages and there was no difference in mean age between groups (U=55, p=0.525; mean ± SD data were 47.9 ± 16.7 for the Normo-vHIT and 52.1 ± 14.9 for the Hypo-vHIT groups). Nevertheless, analysis of the data revealed age-related changes in the composition of the sensory epithelium. As shown in Fig. 3, this was evident in the striola, where significant correlations with age were found for the total number of HCI (CASPR1+) calyces, the total number of calretinin+ calyces, and the number of damaged (CASPR-) calretinin+ calyces. A significant correlation with age was also found for the percentage of CASPR1+ (HCI) cells respect to total HCs (r=-0.476, p=0.022, n=23). In the peripheral areas of the epithelium, most correlations were less robust (data not shown). Nevertheless, a significant effect of age on the total number of HCI (CASPR1+ calyces) was found (r=-0.424, p=0.044, n=23).

**Figure 3.**
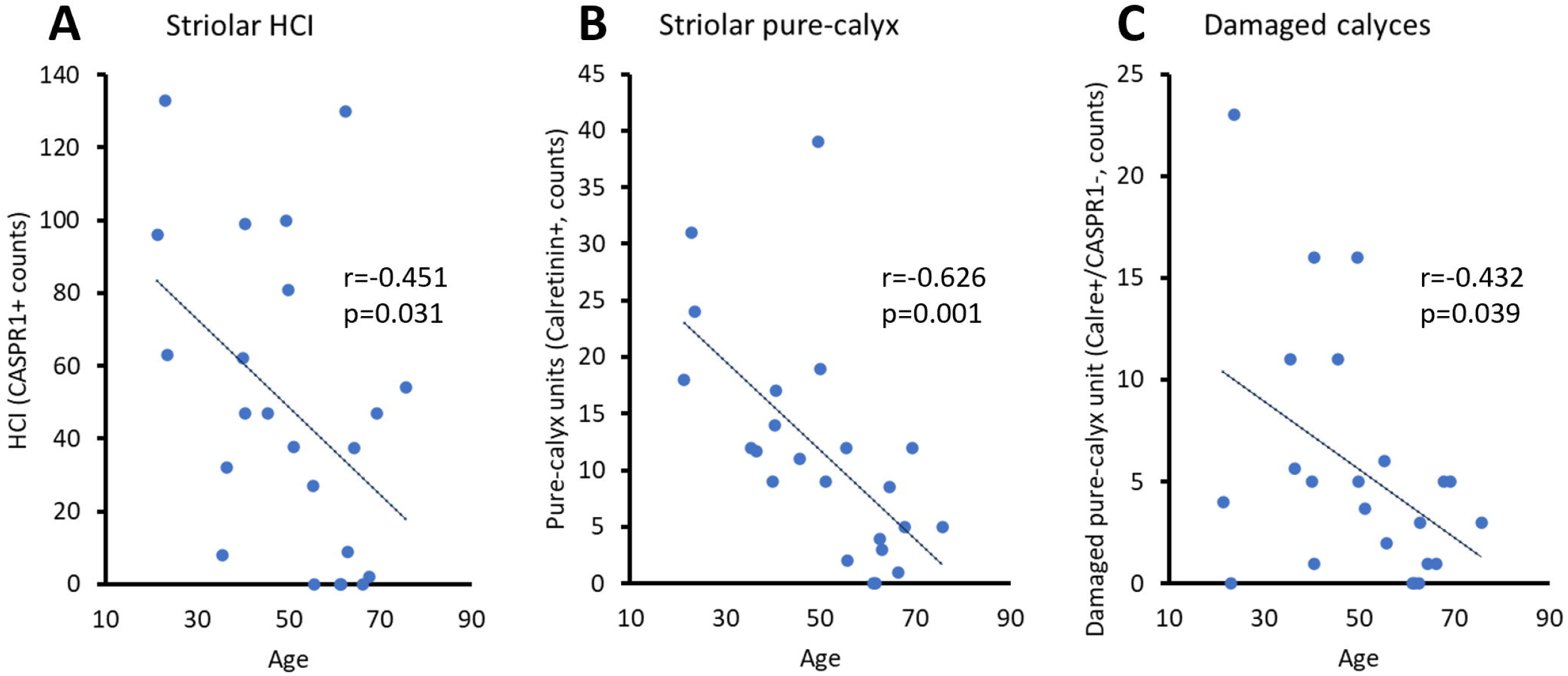
Age-related differences in the structure of the vestibular sensory epithelium of VS patients as revealed by significant correlations with age identified in the striola. **A:** Total number of CASPR1+ calyces. **B:** Total number of calyces of pure-calyx (calretinin+) afferents. **C:** Total number of damaged calyces (CASPR1-) in pure-calyx (calretinin+) afferents. Figures in the panels are Pearson’s correlation coefficient (r) and the corresponding p value (n=23).

As both age and vestibular function group were associated with the histological variables, we used PLSR models to determine the interaction between these factors. Models were generated for the variables that had been found significant in one or the other of the analyses described above: total number of HCI (CASPR1+), total number of calretinin+ calyces, and percent damaged calretinin+ (CASPR1-) calyces respect to total number of calretinin+ calyces. According to the PLSR model results, the total number of HCI decreased with age (t[22 df]=-2.372, p=0.0269) and was higher in Normo-vHIT patients (t[22 df]=2.656, p=0.0144), with no interaction effect (t[22 df]= 0.826, p=0.4178). The total number of calretinin+ calyces decreased with age (t[22 df]=-3.417, p=0.0025), but was not affected by the vHIT-function group (t[22 df]=0.248, p=0.8065) or by the age*group interaction (t[22 df]= −0.140, p=0.8868). In contrast, the percentage of damaged (CASPR1-) calretinin+ calyces was not related to age (t[22 df]=-0.338, p=0.7383), but was significantly lower in Normo-vHIT patients than in Hypo-vHIT patients (t[22 df]=-2.580, p=0.0171); again, the interaction was not significant (t[22 df]=0.828, p=0.4168).

We finally examined the adjusted (Holm correction) significance of the correlations between the histological variables and the quantitative variables from the vHIT tests. The latter included the gains from each semicircular canal in the ipsilateral (VS) side and the contralateral side, as well as the asymmetry between ipsilateral and contralateral sides in these gains. Statistically significant correlations were found for the gain of the ipsilateral lateral canal with the percentage of calretinin+/CASPR1-(damaged pure-calyx) calyces (Fig. 4A), as well as the gain of the ipsilateral anterior canal with 1) the percentage of calretinin+/CASPR1-(damaged pure-calyx) calyces (Fig. 4B), 2) the percentage of striolar HCI (CASPR1+) (R=0.687, p=0.0058, not shown), and 3) the absolute number of striolar HCI (CASPR1+) (Fig. 4C) (Fig. 4). There were no significant correlations with the data from the contralateral side. However, there was a significant correlation (R=0.626, p=0.0319, not shown) between the left anterior – right posterior gain asymmetry and the percentage of striolar HCs showing no calretinin or CASPR1 label, likely corresponding to HCI with dismantled calyceal junction.

**Figure 4.**
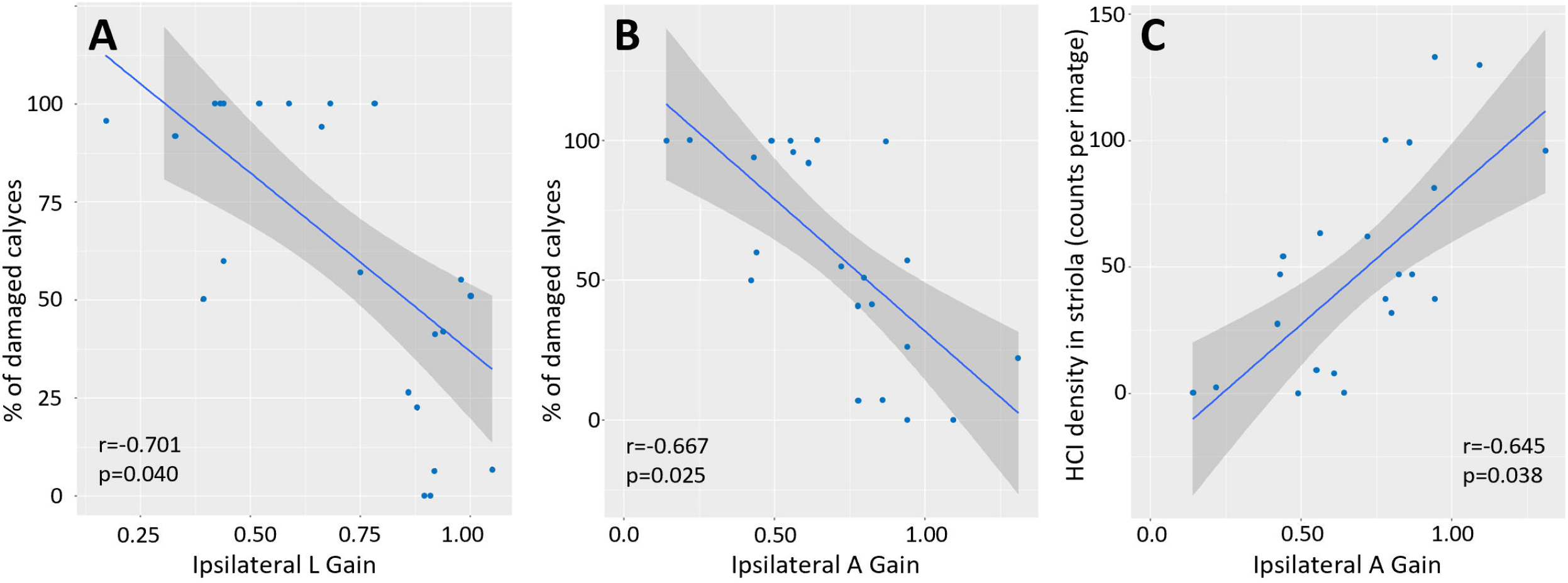
Correlation between functional (vHIT) and histological data. Graphs show individual patient’s data (points) and linear correlation adjustments (line) with 95% confidence intervals (grey shadow). Numbers in each panel are the Pearson’s correlation coefficient (r) and the corresponding p value after the Holm correction (n=23).

## DISCUSSION

The pathological mechanisms responsible for the loss of vestibular function in VS and advanced age remain insufficiently understood. The present study characterizes the pathology of the vestibular sensory epithelium associated with vestibulo-ocular reflex alterations as measured by vHIT in VS patients requiring therapeutic excision of the tumour by a translabyrinthine approach. As these patients had a wide variation in age, the functional and pathological data were evaluated also regarding this factor. The identity of the VS was confirmed by genetic analyses revealing the characteristic loss of function in the *NF2* gene in all cases for which the genetic study was available (18 out of 23).

Several studies have been published reporting on the inner ear pathology in VS cases (Benitez et al., 1967; Hızlı et al., 2016; Moura, 1967; Sans et al., 1996; Ylikoski et al., 1980, 1978), but the use of immunofluorescence labelling is a way of obtaining information related to cellular and molecular features that has been scarcely used in this topic (Wang et al., 2024). In the present study, we have inquired on the impact of VS and age on the density of HC types (I versus II) and on the integrity of the calyceal junction between HCI and the calyx afferents. This junction has been shown to be a sensitive target in animal models of chronic ototoxic stress, in which calyceal junction dismantlement strongly associates with the early evidence of vestibular dysfunction (Borrajo et al., 2025; Greguske et al., 2019; Maroto et al., 2023; Sedó-Cabezón et al., 2015).

Most VS patients show significant vestibular loss, but the extent of this loss is highly variable and does not relate well with tumour size (Balatková et al., 2023; Blödow et al., 2015; Caye-Thomasen et al., 2007; Nadol et al., 1996). In this study, we first aimed to compare the sensory epithelium between patients presenting good vHIT responses with patients suffering significant vHIT response loss. Direct comparison between these groups revealed that Hypo-vHIT patients had significantly lower numbers of both types of sensory cells (HCI and HCII) in both striola and peripheral areas of the sensory organs. This observation strongly supports the hypothesis that sensory epithelium damage contributes to vestibular function loss in VS. Our data match those by Hizli et al. (Hızlı et al., 2016) showing a similar or greater loss of HCI than HCII, and not the finding by Wang et al (Wang et al., 2024) of a greater shortage of HCII in the utricles from VS patients. The HC types were defined by their molecular markers, so the data may also reveal the response of the HCs to the chronic stress associated with the VS. Previous animal research has demonstrated that chronic ototoxic stress causes a reduction in the expression of HC markers including calretinin and MYO7A (Borrajo et al., 2025; Maroto et al., 2023), and loss of the neuronal CASPR1 that characterizes calyces on HCI and has been used here for HCI counts (Borrajo et al., 2025; Greguske et al., 2019; Maroto et al., 2023; Sedó-Cabezón et al., 2015). Therefore, the reduction in HCI and HCII counts may reflect loss of their markers (CASPR1 and calretinin) in addition to the HC loss. In agreement with this hypothesis, the percentage of damaged calyces of pure-calyx afferents (present but without CASPR1) was robustly increased in the Hypo-vHIT group. As pointed out previously (Borrajo et al., 2025) the changes induced by stress in the expression of HC-specific molecular markers adds difficulty to the interpretation of this kind of data. Notwithstanding, the present results provide strong support to the conclusion that, in comparison with the epithelia from patients with more preserved vHIT responses, the epithelia of worse function (Hypo-vHIT) patients contain a higher presence of the molecular pathology previously identified in epithelia submitted to chronic ototoxic stress in animal models, and that, therefore, sensory epithelium stress likely contributes to VS-induced vestibular loss.

Our data also showed an age-related decline in HCI, including a decreasing number of HCI contacted by calyces from pure-calyx afferents. The decrease in the percentage of these pure-calyx (calretinin+) units respect to total HCI (CASPR1+) indicate a greater loss of these units in comparison to all calyces. These data agree with the published evidence of HC loss, and particularly HCI loss with age (Anniko, 1983; Merchant et al., 2000; Rauch, 2001). Importantly, the PLRS models confirmed the effects of age and vestibular function group factors on the histological parameters, and the lack of significant interaction between these factors. These analyses allowed to conclude that HCI decline with age, and that a lower number of HCI units and a higher percentage of damaged calyces associate with a worse vestibular function in VS patients, providing further evidence that VS cause vestibular function loss at least in part through sensory epithelium damage.

We also correlated the histology data with the vHIT measures irrespective of age and function group. After conservative adjustment, significant correlations were found for some parameters with the ipsilateral canal gains, both the lateral and particularly the anterior canals. Thus, larger vHIT gains positively correlate with the number of HCI and the percentage of HCI, suggesting that HCI loss results in a decrease in gain. This is in good agreement with the animal data that indicate that the vestibulo-ocular reflex mainly depends on HCI, rather than HCII, function (Schenberg et al., 2026, 2023). In addition, a higher percentage of damaged pure-calyx calyces associated with lower gains. As this parameter did not correlate with age, this observation supports the conclusion that VS-induced epithelium stress causes dismantlement of the calyceal junction and subsequent loss of the vestibulo-ocular reflex.

One limitation of this study is that vestibular function was only assessed by the vHIT test and did not include other useful tests such as the caloric test or the subjective visual vertical test. Literature data demonstrate that these tests evaluate different aspects of vestibular function and that their combined use results in high sensitivity for diagnosis of VS (Comacchio et al., 2026). However, the present study was not designed to improve or evaluate diagnosis tools and vHIT assessment provided the most efficient way to obtain functional data likely related to the pathology under study, with a focus on the calyx and the calyceal junction.

In summary, we evaluated vestibular function and vestibular epithelium pathology in VS patients of a wide range of ages. The data collected support the hypotheses that 1) the vestibulo-ocular reflex relies more on HCI than HCII; 2) there’s and age-related loss of HCI; 3) VS causes vestibular epithelium damage resulting in vestibular function loss; and 4) the effects of age and VS have an additive, independent, effect on the vestibular epithelium. Importantly, the pathological features associated with the VS-induced effect match those identified in animal models of chronic ototoxicity, which have been demonstrated to be reversible in their early steps. Therefore, the hypothetical removal of the VS while preserving both the labyrinth and vestibular nerve or the inactivation of the tumours’ stressing secretions (e.g., by yet unavailable pharmacological approaches) could eventually lead to partial recovery of the vestibular function of the patient. Likewise, pharmacological interventions providing protection to the vestibular epithelium against the VS-induced stress could mitigate function loss.

## ACKNOWLEDGEMENTS

Funding: Grant 202007-30-31 funded by Fundació La Marató de TV3 and Grant PID2024-155443OB-I00 funded by MICIU/AEI/10.13039/501100011033 and European Regional Development Fund/European Union. MB was supported by the Formación del Profesorado Universitario (FPU) program (Ministerio de Universidades). A.B.G is a Serra-Húnter fellow (Generalitat de Catalunya). The confocal microscopy studies were performed at the Centres Cientifics i Tecnologics de la UB (CCiTUB) of the Universitat de Barcelona. We thank Benjamin Torrejon-Escribano, Irene Boluda, Gemma Casals, and Ignasi Jarne for their expert and technical help. We also thank Fábio Pértille for scientific advice and Francesc Roca-Ribas and Eduard Serra for discussion.

## AUTHOR CONTRIBUTIONS

All authors: conception of the study; design of the work; acquisition, analysis and interpretation of data; revision of the manuscript. MB: visualization; formal analysis. AC: patient assessment; formal analysis; drafting of manuscript. EC: funding acquisition; project administration. EA: patient assessment; supervision. JL: funding acquisition; project administration; supervision; drafting of manuscript.

